# Myosin II activity is required for structural plasticity at the axon initial segment

**DOI:** 10.1101/101584

**Authors:** Mark D Evans, Candida Tufo, Adna S Dumitrescu, Matthew S Grubb

## Abstract

In neurons, axons possess a molecularly defined and highly organised proximal region – the axon initial segment (AIS) – that is a key regulator of both electrical excitability and cellular polarity. Despite existing as a large, dense structure with specialised cytoskeletal architecture, the AIS is surprisingly plastic, with sustained alterations in neuronal activity bringing about significant alterations to its position, length or molecular composition. However, although the upstream activity-dependent signalling pathways that lead to such plasticity have begun to be elucidated, the downstream mechanisms that produce structural changes at the AIS are completely unknown. Here, we use dissociated cultures of rat hippocampus to show that two forms of AIS plasticity in dentate granule cells – long-term relocation, and more rapid shortening – are completely blocked by treatment with blebbistatin, a potent and selective myosin II ATPase inhibitor. These data establish a link between myosin II and AIS function, and suggest that myosin II’s primary role at the structure may be to effect activity-dependent morphological alterations.

## Introduction

Two defining features of neurons – their ability to fire action potentials, and their maintenance of strictly defined cellular polarity – both crucially depend on a molecularly and structurally specialised region of the proximal axon known as the axon initial segment (AIS; Rasband, 2010; Bender & Trussell, 2012). This structure contains a tightly organised and highly regulated arrangement of, amongst others, cytoskeletal components, scaffolding molecules, cell adhesion factors and ion channels, yet is also capable of striking structural plasticity (Grubb & Burrone, 2010a; Grubb *et al.*, 2011). In response to prolonged alterations in levels or patterns of neuronal activity, the AIS can change its location, length and/or its molecular composition, and these changes have been associated with significant alterations in neuronal excitability (Grubb & Burrone, 2010b; Kuba *et al.*, 2010, 2014, 2015, Evans *et al.*, 2013, 2015; Gutzmann *et al.*, 2014; Muir & Kittler, 2014; Chand *et al.*, 2015; Horschitz *et al.*, 2015; Wefelmeyer *et al.*, 2015; Dumitrescu *et al.*, 2016). Structural change at the AIS may therefore be part of a repertoire of plastic mechanisms by which neurons can alter their function in response to perturbations in their ongoing electrical activity.

However, although the AIS is known to be structurally plastic in multiple ways, the cellular mechanisms responsible for these activity-dependent structural changes are almost entirely unknown. Some headway has been made in elucidating the upstream signalling pathways necessary for AIS plasticity. In all cell types tested this is dependent upon calcium entry through L-type Ca_V_1 voltage-gated calcium channels, and in hippocampal dentate granule cells both long-term (48 h) AIS relocation and more rapid (3 h) AIS shortening are mediated by the activity of the calcium-dependent phosphatase calcineurin (Evans *et al.*, 2013, 2015; Chand *et al.*, 2015). But which cellular events downstream of calcineurin activity are necessary to produce structural changes at the AIS? Here, cytoskeletal components of the AIS are prime suspects. Microtubule− and actin-based dynamics are key players driving diverse forms of morphological change in neurons, and the AIS is known to possess uniquely organised structures of both microtubules and actin filaments (Palay *et al.*, 1968; Watanabe *et al.*, 2012; Xu *et al.*, 2013; Jones *et al.*, 2014; D’Este *et al.*, 2015; Ganguly *et al.*, 2015; Leterrier *et al.*, 2015). Indeed, these cytoskeletal specialisations have long been proposed to play a role in altering AIS structure (Palay *et al.*, 1968), and recent data show that altering microtubule stability can affect AIS location in hippocampal neurons (Hatch *et al.*, 2017). Axons are also rich in actin-associated myosin motor proteins, with non-muscle myosin II playing crucial roles in growth cone motility, and myosin Va and myosin VI controlling the AIS-based filtering of dendritically− or axonally-targeted proteins, respectively (Arnold & Gallo, 2014). To date, though, myosins have not been implicated in the development, maintentance or plasticity of AIS structure *per se*. Here we demonstrate just such a role, using pharmacological manipulations in dissociated hippocampal cultures to show that the activity of myosin II is necessary for multiple forms of structural AIS plasticity.

## Methods

### Dissociated hippocampal culture

Humane killing for tissue collection conformed to local King’s College London ethical approval under the UK Supplementary Code of Practice, The Humane Killing of Animals under Schedule 1 to the Animals (Scientific Procedures) Act 1986. Hippocampi were rapidly dissected from embryonic day (E18) Wistar rat embryos (Charles River) of either sex in ice-cold Hank’s balanced salt solution (HBSS). Tissue was trypsin digested (Worthington, 0.5 mg/ml; 15 min at 37°C), then triturated by repeatedly pipetting the cells using fire-polished Pasteur pipettes, and finally plated on 13 mm coverslips (45,000 cells/coverslip; VWR) coated with poly-l-lysine (50 μg/ml, Sigma) and laminin (40 μg/ml). Cells were incubated at 37°C with 5% CO_2_ in Neurobasal medium containing 1% B27, 1% foetal calf serum and 500 μM Glutamax. At 4 days *in vitro* (DIV) half the media was changed with Neurobasal plus 2% B27 and 500 μM Glutamax. At 7 DIV media was topped up to 1 ml with fresh Neurobasal plus 2% B27 and 500 μM Glutamax. All experiments were carried out between 10–12 DIV. Unless otherwise stated, all cell culture reagents were obtained from Invitrogen.

### Transfections

EGFP–C1 nuclear factor of activated T-cells 3 (NFAT3; referred to as NFAT–GFP) was obtained from Addgene (plasmid 10961; deposited by Toren Finkel, National Heart, Lung and Blood Institute, Bethesda, MD, USA; Ichida & Finkel, 2001). Neuronal cultures were sparsely transfected with NFAT– GFP at 7 DIV using Lipofectamine 2000 (Invitrogen; 0.5 µg DNA and 0.5 µl lipofectamine per well, in 1 ml media; 10 min at 37°C).

### Neuronal treatments

All treatments were performed at 10 DIV. Pharmacological agents, (±)-blebbistatin (Abcam; ab120425) and dynasore (Abcam; ab120192), were made up in DMSO at 1000-fold stock concentrations. They were subsequently added to culture media at previously described effective working concentrations (Newton *et al.*, 2006; Kollins *et al.*, 2009) at least 30 min before control or depolarising treatment. Cultures were depolarised by adding +10 mM or +15 mM KCl from 1M stock, or treated with +10 mM or +15 mM NaCl as osmolarity controls.

### Immunocytochemistry

Cultures were fixed in 4 % paraformaldehyde (TAAB Laboratories; in 3 % sucrose, 60 mM PIPES, 25 mM HEPES, 5 mM EGTA, and 1 mM MgCl_2_) for 20 min at room temperature. Cells were permeabilised for 5 min with 0.25 % Triton X-100 (Sigma) before blocking for 1 h in 10 % goat serum (GS; Sigma). Coverslips were then placed in primary antibody solution, in PBS plus 2 % GS, at relevant concentrations: mouse monoclonal anti-ankyrin-G (Neuromab clone N106/36; 1:500) and rabbit polyclonal anti-prox1 (Sigma P1724; 1:1000) for 90 min. After washing in PBS, coverslips were placed in relevant secondary antibody solution (Invitrogen Alexa Fluor-conjugated antibodies in 2% GS; 1:1000) for an additional hour, before further washing and mounting in MOWIOL (Calbiochem).

### Imaging and image analysis

All imaging and subsequent analyses were performed blind to experimental group. For AIS imaging and quantification, prox1-positive dentate granule cells (DGCs) with ankyrin-G-positive AISs of obvious somatic origin were visualized under epifluorescence and imaged using a laser-scanning confocal microscope (Carl Zeiss LSM 710). Neurons were imaged with appropriate excitation and emission filters, with the pinhole set to 1 AU and using a 40× oil-immersion objective (Carl Zeiss). Laser power and gain settings were adjusted to prevent signal saturation. Images were taken with 3× zoom, 512 × 512 pixels (0.138 μm/pixel), and in *z*-stacks with 0.5 μm steps.

Stacks were flattened into single maximum intensity projections and imported into MATLAB (MathWorks) for analysis using custom-written functions (Evans *et al.*, 2015; freely available at www.mathworks.com/matlabcentral/fileexchange/28181-ais-quantification). We drew a line profile starting at the soma that extended down the axon, through and past the AIS. At each pixel along this profile, fluorescence intensity values were averaged over a 3 × 3 pixel square centered on the pixel of interest. Averaged profiles were then smoothed using a 40-point (∼5 μm) sliding mean and normalized between 1 (maximum smoothed fluorescence, location of the AIS max position) and 0 (minimum smoothed fluorescence). AIS start and end positions were obtained at the proximal and distal axonal positions, respectively, at which the normalized and smoothed profile declined to 0.33.

For 48 h-treated cells, an AIS movement index (AMI; Evans *et al.*, 2013) was calculated for each neuron subjected to both drug and depolarisation using its own AIS max position (*a*) and the mean AIS max position for other treatment groups [drug + 10 mM NaCl 
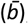
; solvent + 10 mM KCl 
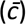
; and solvent + 10 mM NaCl 
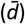
] as follows:

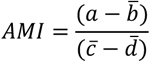

A mean AMI of 1 describes an experiment in which AISs moved as far in drug as in control, whereas an AMI of 0 indicates that AIS relocation has been totally blocked.

The distribution of NFAT–GFP (Evans *et al.*, 2013) was assessed in prox1-positive neurons by tracing separate nuclear and cytoplasmic regions within a single-plane confocal image (ImageJ, NIH), and measuring the mean grey value for both nuclear and cytoplasmic NFAT–GFP. Then,

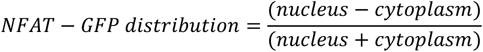

## Statistics

Data were analysed with Prism software (GraphPad Software). Datasets were assessed for Normality using a D’Agostino and Pearson omnibus test, and were analysed using parametric or nonparametric tests accordingly. All tests were two-tailed, with α = 0.05.

## Results

We tested the involvement of myosin II in structural AIS plasticity by using the potent and selective non-muscle myosin II ATPase inhibitor blebbistatin (50 µM; Straight *et al.*, 2003; Limouze *et al.*, 2004; Kollins *et al.*, 2009) during neuronal depolarisation in dissociated cultures of rat hippocampus. We first investigated the slow distal relocation of the AIS produced by +10 mM KCl depolarisation over 48 h (Grubb & Burrone, 2010b; Evans *et al.*, 2013; Muir & Kittler, 2014; Horschitz *et al.*, 2015), focusing on dentate granule cells (DGCs) revealed by immunocytochemical label for the transcription factor prox1 (Williams *et al.*, 2011; Evans *et al.*, 2013; Lee *et al.*, 2013). AIS position was quantified based on immunolabel for the scaffolding molecule ankyrin-G (AnkG). In the presence of the DMSO solvent control, 48 h depolarisation produced the expected ~ 9 µm distal relocation of the entire AIS structure, with the start, max, and end AIS positions all shifting significantly down the axon away from the soma (Fig. 1; Start position mean ± SEM: +10 mM NaCl 1.73 ±0.50 µm, +10 mM KCl 10.06 ± 0.96 µm, Tukey post-test after 2-way ANOVA on ranks, p < 0.0001; Max position: NaCl 9.97 ± 0.86 µm, KCl 19.12 ± 1.17 µm; p < 0.0001; End position: NaCl 22.40 ± 0.81 µm, KCl 31.48 ± 1.36 µm, p < 0.0001). In the presence of 50 µm blebbistatin, however, this activity-dependent relocation of the AIS was blocked completely. AIS start, max and end positions were no different from those under non-depolarised conditions (Fig. 1; Start position mean ± SEM: +10 mM NaCl 2.36 ±0.49 µm, +10 mM KCl 2.09 ± 0.45 µm, Tukey post-test after 2-way ANOVA on ranks, p = 0.99; Max position: NaCl 10.00 ± 0.68 µm, KCl 9.62 ± 0.70 µm; p > 0.99; End position: NaCl 22.24 ± 0.78 µm, KCl 23.79 ± 0.74 µm, p = 0.58), producing an AIS movement index (AMI; see Methods) that was not significantly different from zero (mean ± SEM −0.041 ± 0.076; 1-sample t-test vs 1, p < 0.0001; vs 0, p = 0.59).

**Fig 1:**
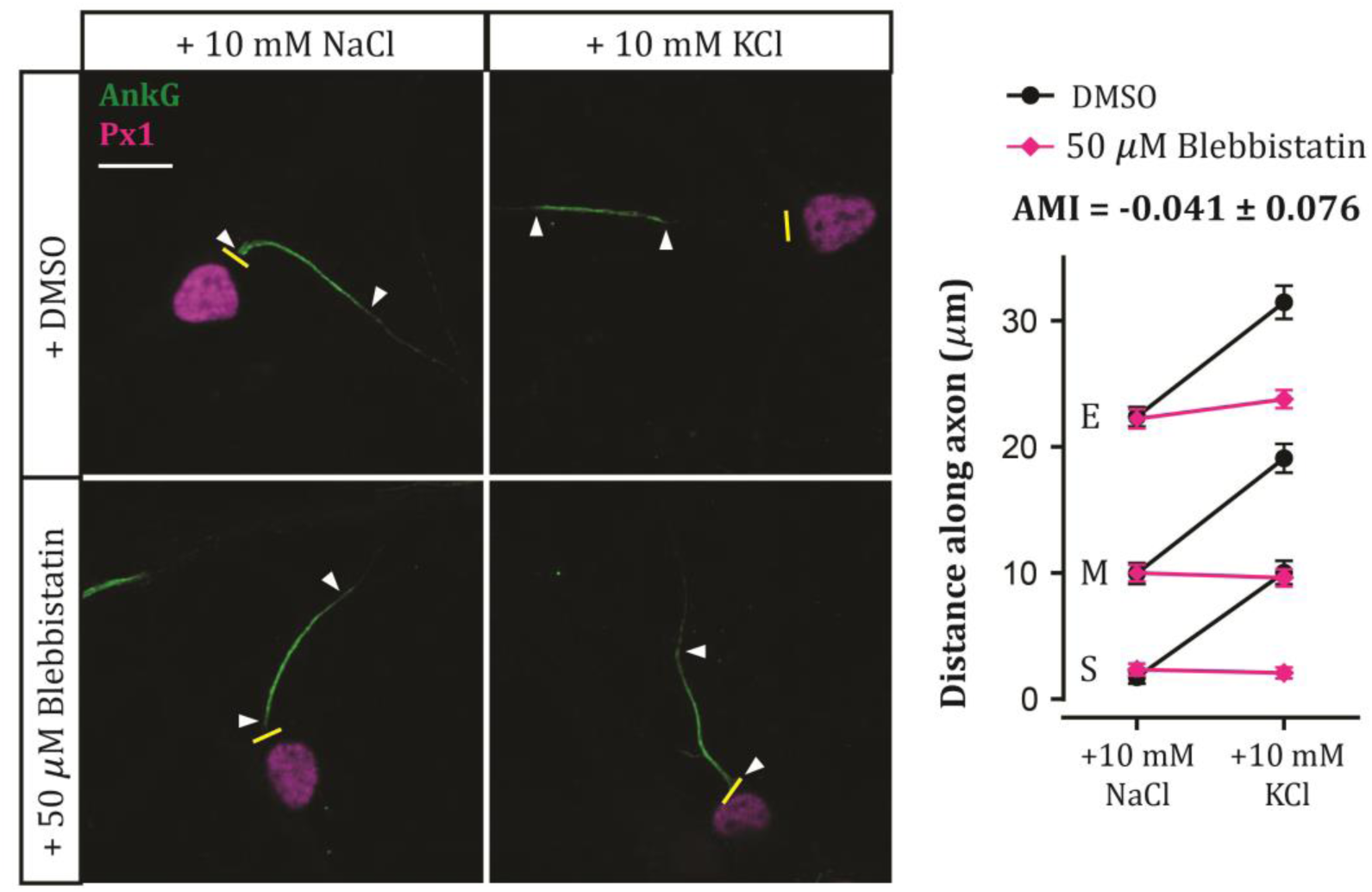
Myosin II is necessary for 48 h AIS relocation. Example maximum intensity projection images (left) of neurons labelled for ankyrin-G (AnkG) and prox1 (Px1) following 48 h NaCl or KCl treatment in the presence of DMSO or 50 μM blebbistatin. Yellow lines, axon start; white arrowheads, AIS start and end positions; scale bar, 10 μm. Plot (right) shows mean ± SEM of AIS start (S), max (M) and end (E) position for each treatment group. AMI, AIS movement index (see Methods).

Myosin II is therefore necessary for slow activity-dependent AIS relocation over several days. But structural plasticity at the AIS can also occur much more rapidly, with just 3 h depolarisation capable of producing a significant decrease in the structure’s length (Evans *et al.*, 2015). We asked whether myosin II is also necessary for such rapid AIS shortening, by treating our neurons with 50 µM blebbistatin during 3 h depolarisation with +15 mM KCl. As expected, we found a significant decrease in AIS length after 3 h KCl treatment in DMSO (Fig. 2; mean ± SEM, +15 mM NaCl 19.80 ± 0.59 µm, +15 mM KCl 16.47 ± 0.96 µm; Tukey post-test after 2-way ANOVA, p = 0.0054). This activity-dependent AIS shortening was entirely blocked, however, in the presence of 50 µM blebbistatin (Fig. 2; mean ± SEM, +15 mM NaCl 19.46 ± 0.57 µm, +15 mM KCl 20.29 ± 0.62 µm; Tukey post-test after 2-way ANOVA, p = 0.84).

**Fig 2:**
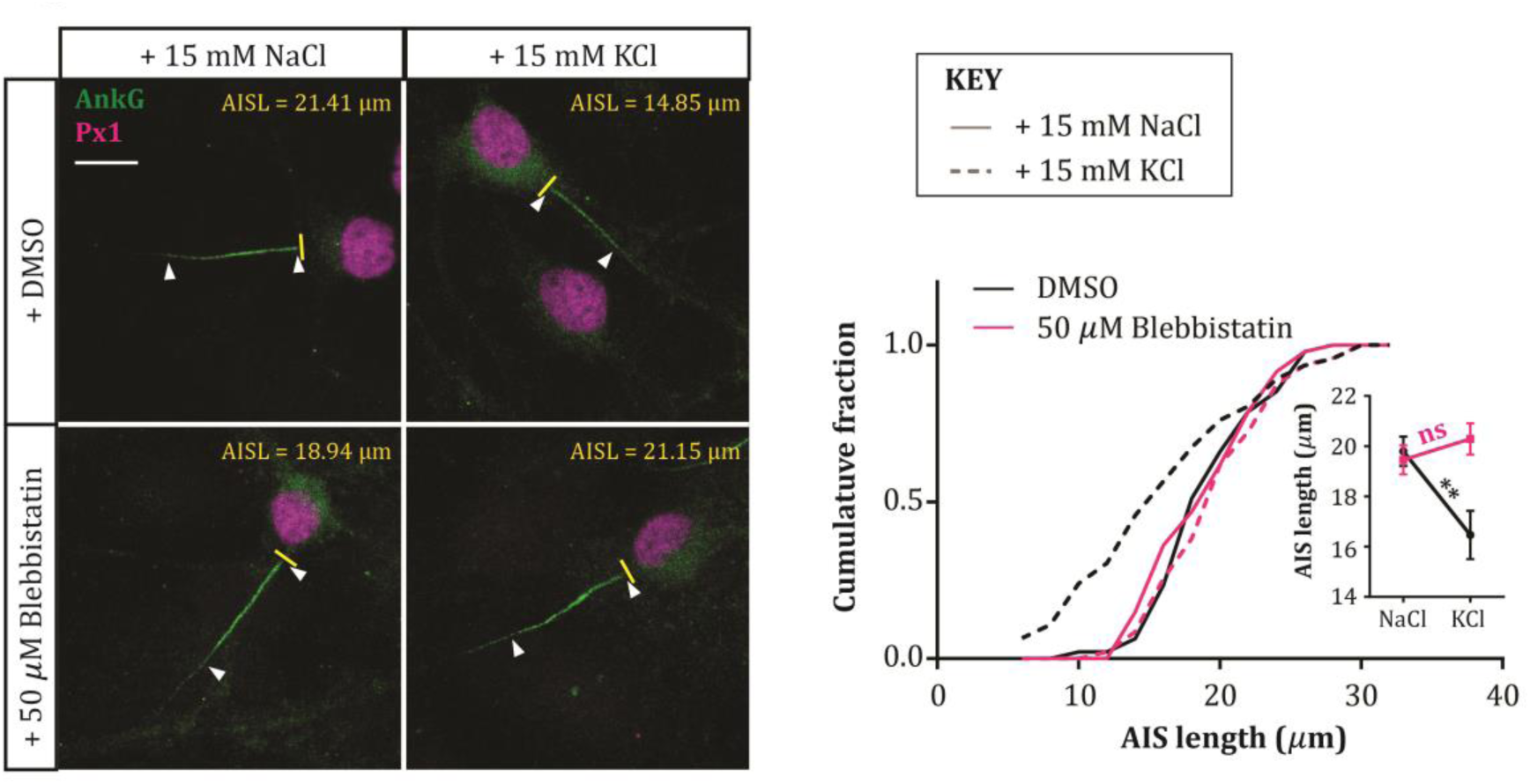
Myosin II is necessary for 3 h AIS shortening. Example maximum intensity projection images (left) of neurons labelled for ankyrin-G (AnkG) and prox1 (Px1) following 3 hr NaCl or KCl treatment in the presence of DMSO or 50 μM blebbistatin. Yellow lines, axon start; white arrowheads, AIS start and end positions; AISL, AIS length for each image; scale bar, 10 μm. Plot (right) shows cumulative fraction and (inset) mean ± SEM of AIS length. Two-way ANOVA with Tukey’s multiple comparison test; ** p = 0.0054; ns, non-significant.

In hippocampal DGCs, both slower AIS relocation and rapid AIS shortening are known to depend on signalling through L-type Ca_V_1 voltage-gated calcium channels and the calcium-dependent phosphatase calcineurin (Evans *et al.*, 2013, 2015). Could blebbistatin be preventing AIS plasticity by somehow inhibiting these upstream signalling events? We tested this possibility by transfecting our cells with the calcineurin-dependent probe NFAT-GFP, which is localised to the cytoplasm under baseline conditions but is rapidly translocated to the nucleus upon calcineurin activation (Graef *et al.*, 1999). Indeed, here 3 h depolarisation with +15 mM KCl in DMSO produced clear nuclear translocation of the NFAT-GFP probe (Fig. 3; NFAT distribution mean ± SEM, +15 mM NaCl − 0.49 ± 0.02, +15 mM KCl 0.64 ± 0.02; Tukey post-test after 2-way ANOVA, p < 0.0001). This nuclear translocation was also clearly observed in the presence of 50 µM blebbistatin, albeit to a slightly reduced extent (Fig. 3; mean ± SEM, +15 mM NaCl − 0.42 ± 0.03, +15 mM KCl 0.41 ± 0.03; Tukey post-test after 2-way ANOVA, p < 0.0001; Tukey post-test for DMSO vs blebbistatin in KCl, p < 0.0001). Myosin II is therefore required for structural AIS plasticity at some point downstream of calcium-dependent calcineurin signalling.

**Fig 3:**
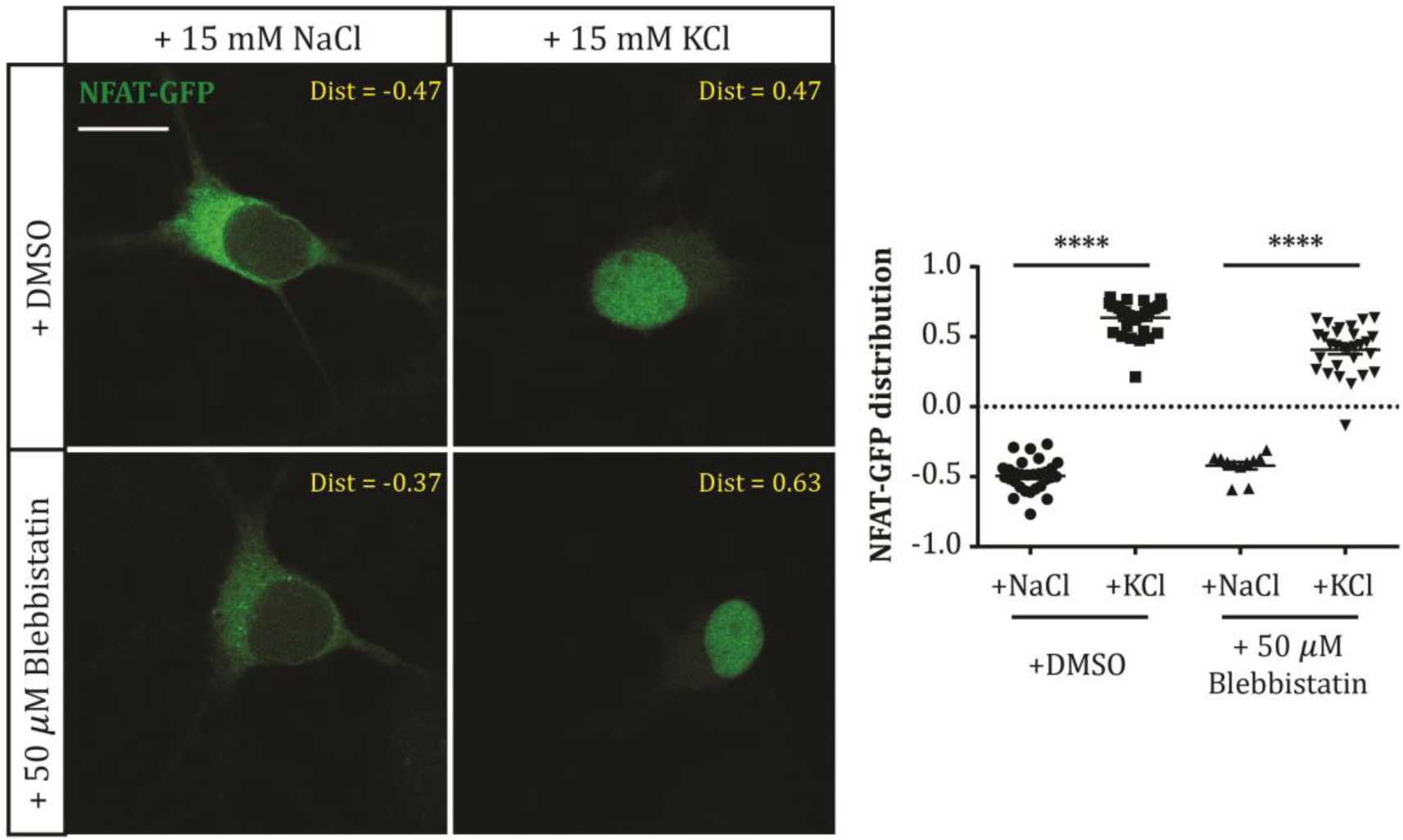
Myosin II inhibition does not block activity-dependent calcineurin signalling. Example single-plane images (left) of NFAT-GFP-expressing DGCs after treatment with NaCl or KCl in the presence of DMSO or 50 μM blebbistatin. Dist, nucleus:cytoplasm distribution for each image; scale bar, 10 μm. Plot (right) shows NFAT-GFP nucleus: cytoplasm distribution (see Methods) in each treatment group. Each symbol represents one cell; lines show mean ± SEM. ****, 2-way ANOVA with Tukey’s multiple comparison test, p < 0.0001.

One cellular process known to depend on both calcineurin and myosin II in neurons is bulk endocytosis (Evans & Cousin, 2007; Flores *et al.*, 2014). Could this *en masse* removal of cell membrane be a crucial mechanism in mediating AIS plasticity? We tested this hypothesis by blocking another key component in the bulk endocytosis process – dynamin (Clayton *et al.*, 2009; Nguyen *et al.*, 2012; Gormal *et al.*, 2015) – using the dynamin inhibitor dynasore (80 µM; Macia *et al.*, 2006; Newton *et al.*, 2006). However, although dynasore had a depolarisation-independent effect on AIS length, producing a ~2.5 µm decrease after both 3 h +15 mM NaCl and KCl treatments (Fig. 4; DMSO mean ± SEM, +15 mM NaCl 20.65 ± 0.76 µm, KCl 14.90 ± 0.56 µm; dynasore, NaCl 18.17 ± 0.62 µm, KCl 12.09 ± 0.66 µm; effect of drug in 2-way ANOVA, F_1,135_ = 16.44, p < 0.0001), it had no influence whatsoever on activity-dependent rapid AIS shortening (Fig. 4; effect of drug x treatment interaction in 2-way ANOVA, F_1,135_ = 0.06, p = 0.80; NaCl vs KCl in DMSO, and NaCl vs KCl in dynasore, Tukey post-test after 2-way ANOVA, both p < 0.0001).

**Fig 4:**
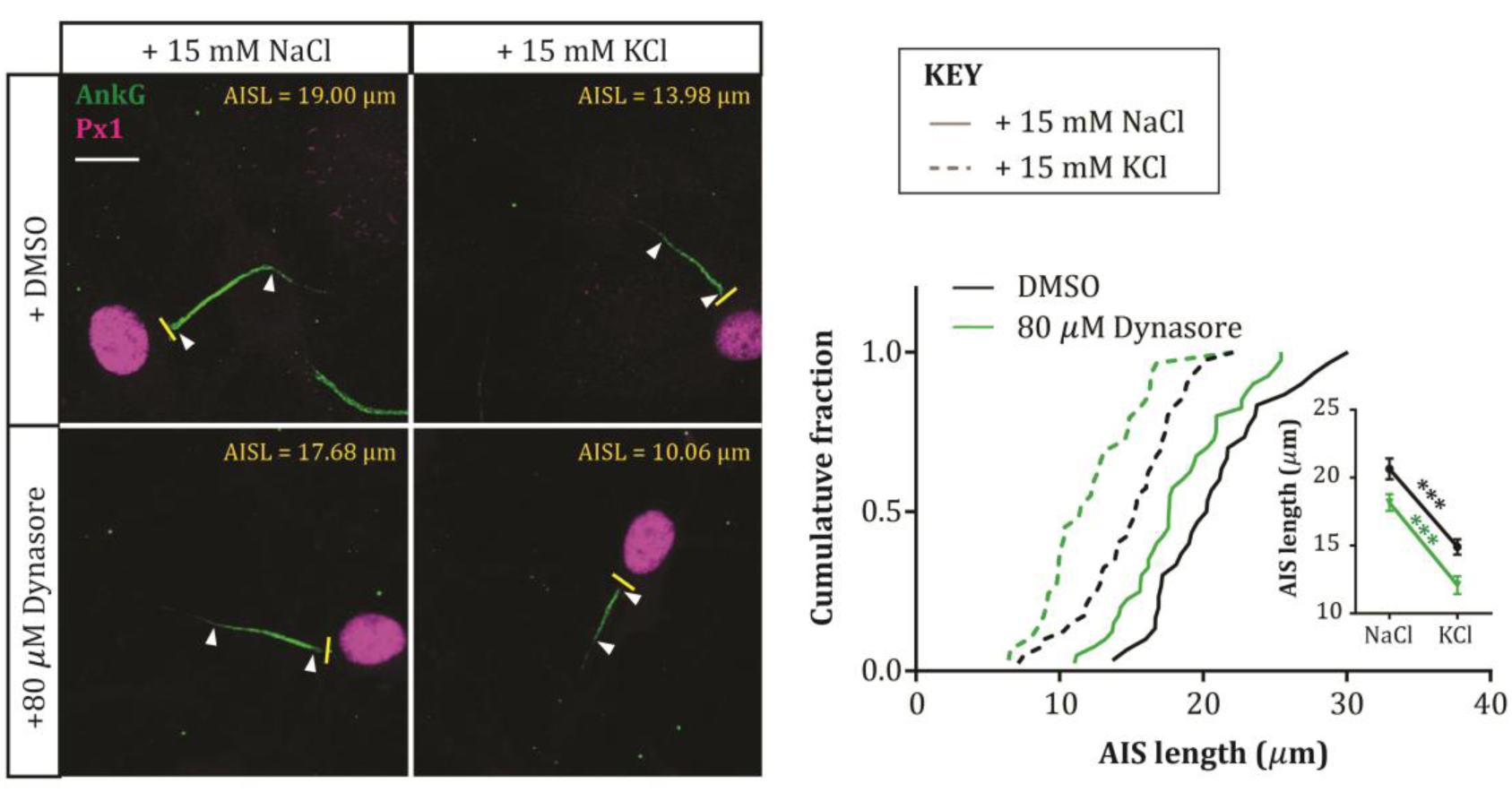
AIS shortening is not dependent upon endocytosis. Example maximum intensity projection images (left) of neurons labelled for ankyrin-G (AnkG) and prox1 (Px1) following 3 hr NaCl or KCl treatment in the presence of DMSO or 80 μM dynasore. Yellow lines, axon start; white arrowheads, AIS start and end positions; AISL, AIS length for each image; scale bar, 10 μm. Plot (right) shows cumulative fraction and (inset) mean ± SEM of AIS length. Two-way ANOVA with Tukey’s multiple comparison test; ***, p < 0.0001.

## Discussion

We show here that structural plasticity at the AIS is blocked by treatment with blebbistatin, a selective myosin II ATPase inhibitor. Furthermore, we show that the effect of myosin II inhibition cannot be explained by upstream effects on calcineurin signalling, or by involvement in bulk endocytosis.

How might myosin II be acting to produce activity-dependent structural changes at the AIS? Although our current data cannot formally rule out the possibility that myosin II plays an indirect role in AIS plasticity by acting elsewhere in dentate granule cells, or in the hippocampal network, the most parsimonious explanation for its requirement in AIS plasticity is that it is directly involved in the cytoskeletal arrangements that produce AIS shortening and relocation. This would certainly be in keeping with myosin II’s well-characterised roles in stimulus-induced structural changes and activity-dependent plasticity elsewhere in the neuron. In developing distal axons, myosin II has well-characterised roles in growth cone motility (Vallee *et al.*, 2009) and sema3A-induced axon retraction (Gallo, 2006; Arnold & Gallo, 2014). In dendrites it is necessary for spine maturation (Ryu *et al.*, 2006; Rubio *et al.*, 2011; Koskinen *et al.*, 2014), while distinct myosin II isoforms are also critical for the normal maturation of specific dendritic compartments and pathway-specific synaptic function (Ozkan *et al.*, 2015). Myosin II activity is also crucial for the stability of long-term potentiation (Rex *et al.*, 2010).

Recent evidence that myosin II can interact directly with the master AIS scaffolding molecule ankyrin-G (Dash *et al.*, 2016) suggests that it could be an integral part of the AIS’s sub-membraneous structure, and may be in an ideal position to affect molecular change in response to alterations in electrical activity. The possibility that myosin II might be specifically involved in AIS plasticity, rather than in maintenance of AIS structure *per se*, is also underlined by the complete lack of effect on AIS length or position when its activity is blocked under baseline activity conditions, even for 48 h (Fig. 1). This suggests that myosin II could be a specialised AIS component, called into action only when structural alterations are required in response to chronic alterations in ongoing activity.

But how exactly might myosin II produce structural changes at the AIS? This is most likely to depend on interactions with the specialised actin cytoskeleton in the proximal axon which not only organises – like the rest of the axon - into stable, regularly-spaced transverse rings linked by longitudinally-arranged spectrin molecules, but also contains distinct ‘patches’ where actin is densely organised into non-parallel meshworks, as well as deeper actin ‘hotspots’, which may generate a more dynamic actin cytoskeletal network (Watanabe *et al.*, 2012; Xu *et al.*, 2013; D’Este *et al.*, 2015; Ganguly *et al.*, 2015; Leterrier *et al.*, 2015). Interestingly, the mechanism of action of myosin II has been shown to depend on the organisation of the actin filaments with which it interacts – while parallel actin filaments are more likely to be stabilised by myosin II, the same motor protein can contract and disassemble antiparallel filaments (Reymann *et al.*, 2012). At the AIS, myosin II’s role in active actin depolymerisation may therefore be a crucial step in breaking down the tight relationship between the ankyrin-G/βIV-spectrin scaffold and the actin cytoskeleton (Xu *et al.*, 2013; Leterrier *et al.*, 2015), especially at non-parallel actin patches or hotspots. This may then allow removal of AIS components in rapid activity-dependent shortening, to be followed by subsequent AIS relocation. Alternatively, or in addition, a more canonical myosin II contractile motor function might be involved in the mechanics of AIS structural plasticity, perhaps via interactions with axonal actin ring structures. No doubt super-resolution microscopy investigating the interactions between myosin II and actin, both at rest and during conditions of elevated neuronal activity, will soon shed more light on the precise molecular mechanisms of AIS plasticity.

Upstream factors known to mediate activity-dependent structural change at the AIS include L-type Ca_V_1 calcium channels and the calcium-activated phosphatase calcineurin (Evans *et al.*, 2013, 2015; Chand *et al.*, 2015). How might these signalling pathways be linked to myosin II activity? Discounting any direct effects of myosin II on calcineurin signalling itself (Fig. 3), or any role for calcineurin− and myosin II-dependent bulk endocytosis in AIS plasticity (Fig. 4), prime suspects include direct calcineurin activation of the cofilin phosphatase slingshot (Wen *et al.*, 2007; Yuen & Yan, 2009), or indirect modulation of actin-myosin II interactions via calcineurin-dependent regulation of protein synthesis (Graef *et al.*, 1999; Flavell *et al.*, 2006; Qiu & Ghosh, 2008; Li *et al.*, 2009), and these remain promising targets for future investigation. Finally, uncovering a specific molecular player in the process of AIS plasticity may also shed light on neuronal strategies for integrating multiple mechanisms of activity-dependent change (Turrigiano, 2011). While calcineurin signalling is known to be involved in multiple, sometimes opposing plastic processes (Evans *et al.*, 2015), blocking AIS changes alone via specific inhibition of myosin II might allow the functional effects of structural AIS plasticity to be studied in effective isolation.

## Acknowledgements

This work was supported by a Wellcome Trust Career Development Fellowship to MSG (088301), and Medical Research Council 4-year PhD studentships to MDE and ASD. We thank Annisa Chand for assistance with cultures, Caroline Formstone for sharing reagents, and Phillip Gordon-Weeks for comments on the manuscript.

## Author contributions

MDE, ASD and MSG designed experiments; all authors performed experiments; MDE, CT and MSG analysed data; MDE and MSG wrote the paper.

